# Electron microscopic analysis of the influence of iPSC-derived motor neurons on bioengineered human skeletal muscle tissues

**DOI:** 10.1101/2023.03.03.530083

**Authors:** Christine T. Nguyen, Carolina Chávez-Madero, Erik Jacques, Brennen Musgrave, Ting Yin, Kejzi Saraci, Penney M. Gilbert, Bryan A. Stewart

**Affiliations:** Department of Biology, University of Toronto Mississauga, Mississauga, ON, L5L 1C6, Canada; Department of Cell & Systems Biology, University of Toronto, Toronto, ON, M5S 3G5, Canada; Institute of Biomedical Engineering, University of Toronto, Toronto, ON, M5S 3G9, Canada; Donnelly Centre, University of Toronto, Toronto, ON, M5S 3E1, Canada

## Abstract

3D bioengineered skeletal muscle macrotissues are increasingly important for studies of cell biology and development of therapeutics. Tissues derived from immortalized cells obtained from patient samples or from stem cells can be co-cultured with motor-neurons to create models of human neuromuscular junctions in culture. In this study, we present foundational work on 3D cultured muscle ultrastructure, with and without motor neurons, which is enabled by the development of a new co-culture platform. Our results show that tissues from Duchenne muscular dystrophy patients are poorly organized compared to tissues grown from healthy donor and that the presence of motor neurons invariably improves sarcomere organization. Electron micrographs show that in the presence of motor neurons, filament directionality, banding patterns, z-disc continuity and appearance of presumptive SSR and T-tubule profiles all improve in healthy, DMD and iPSC derived muscle tissue. Further work to identify the underlying defects of DMD tissue disorganization and the trophic mechanisms by which motor neurons support muscle are likely to yield potential new therapeutic approaches for treating patients suffering from Duchenne muscular dystrophy.

## Introduction

The advent of improved cell culture techniques, in particular the establishment of patient-derived cell lines and the increasing use of pluripotent stem cells (PSCs), have enhanced our ability to use tissues grown in culture, as platforms to study basic cell biology and screen therapeutic agents for disease treatments. Our research is focused on muscle biology, and we have developed 3D muscle culturing techniques to grow human muscle microtissues, which we have demonstrated provides advantages over traditional 2D cultures (Afshar Bakooshli et al., 2019; Ebrahimi et al., 2021; Moyle et al., 2022; Nguyen et al., 2021). Further, we are using our platform to investigate Duchenne muscular dystrophy (DMD) and identify potential therapeutic treatments (Afshar Bakooshli et al., 2019; Ebrahimi et al., 2021; Nguyen et al., 2021). While there is considerable effort to develop gene-based (Zhang et al., 2021) and cell-based therapies (Galli et al., 2021) for DMD(Mollanoori et al., 2021; Wilton-Clark & Yokota, 2023), such treatments face a long developmental pipeline, and not all patients are eligible. While conventional therapeutic options remain limited (Kracht et al., 2022), we believe that concomitant development of drug therapies that target root physiological perturbations have the potential to improve DMD symptoms and in the long run will complement other therapeutic interventions.

Recently, we generated functional human neuromuscular systems in 3D culture (Bakooshli et al., 2019)and we have compared the physiology and sarcomeric structure of tissues derived from healthy and DMD patients (Ebrahimi et al., 2021; Nguyen et al., 2021). Importantly, we have shown that 3D culturing better matches data obtained from human muscle biopsy than does 2D cultures (Nguyen et al., 2021). Further, we have shown proof in principle that some of the electrophysiological hallmarks of DMD detectable in 3D culture can be improved by known therapeutic treatments, such as polaxamer 180.

In addition, using immortalized cell lines, we have shown at the ultrastructural level that, in the absence of motor neurons (MNs), sarcomeres in DMD derived tissue are disorganized compared to tissue derived from healthy donors. This is an important observation because it is impossible to study DMD muscle *in vivo* or from biopsy without the influence of motor neurons. In our system, we can decouple the cellular components of the neuromuscular junction. To date, we have demonstrated that a β1-integrin activating antibody, improves sarcomere organization, contractile apparatus maturation, and stability, in the absence of neurons (Ebrahimi et al., 2021). The aim of the present study is to build upon our previous ultrastructure work to ask if the presence of motor neurons *per se* alters the ultrastructural development of 3D grown human muscle, using cells obtained from DMD and healthy donors. We utilize an established co-culture method to grow muscle tissue in the presence or absence of motor neurons clusters (Bakooshli et al., 2019), and herein present a new co-culture platform that offers superior support of maturing neuromuscular tissues. Most of these studies use tissue from immortalized cells, but for the first time, we show the ultrastructure of 3D muscle grown from human induced pluripotent stem cells (iPSCs).

Our observations show unequivocally that the presence of motor neurons improves the ultrastructural organization of muscle tissue. Parallel studies on synaptic connections and organization are ongoing.

## Materials and Methods

### Fabrication of MyoFMS culture platform

Fabrication of the culture platform MyoFMS was modelled after a previously developed protocol(Iuliano et al., 2020). More specifically, SLA laser-based 3D printing was performed using a Form3 (Formlabs, USA) and resin clear V4 (RS-F2-GPCL-04); printing parameters were set to Adaptive Layer Thickness. After printing, the resulting 3D piece was cleaned and post-processed as described by the manufacturer. Next, a treatment of trichloro (1H, 1H, 2H, 2H-perfluorooctyl)silane (Millipore Sigma) was applied to the surface of the 3D piece to make it inert and to facilitate the next step of casting. To do this, the 3D printed part was plasma-treated for 1 minute in a plasma chamber (Harrick Plasma); and subsequently, it was placed in a vacuum chamber, exposed to an open container with 250 μL of the trichloro(1H, 1H, 2H, 2H-perfluorooctyl)silane and vacuum was applied for 15 minutes to ensure chemical deposition on the piece. The silane treatment was incubated overnight. The next day, the piece was placed in the oven for 1 hour at 70°C and then washed with isopropanol.

An Ecoflex 00–30 (Smooth-On Inc., PA)-based device was used as a master mold. To generate this master mold from the 3D printed piece, we followed the manufacturer’s protocol. Specifically, Ecoflex parts A:B were mixed in equal ratios on the 3D piece, and a vacuum chamber was used for degassing the material. This was then left to crosslink at room temperature for 4 hours, at which point the Ecoflex was solid and was then carefully separated from the 3D printed piece. Ecoflex molds were exposed to the silane treatment exactly as described above for the 3D-printed part. The Ecoflex mold was then used to generate the polydimethylsiloxane (PDMS) tissue culture platform. No additional coating was needed prior to pouring the liquid PDMS (15:1 w/w monomer:curing agent) onto the Ecoflex mold. The mold coated with liquid PDMS was degassed for 1 hour followed by overnight crosslinking at 60°C. The resulting PDMS was then slowly and carefully released from the Ecoflex at room temperature, and the PDMS-based MyoFMS culture platform was then autoclaved in preparation for implementation in sterile tissue culture applications.

### Human immortalized myoblast cell line maintenance

The human immortalized myoblast cell lines used in this study (AB1190 and AB1071) were obtained from the Institut de Myologie in Paris, France (Mamchaoui et al., 2011)through a material transfer agreement. In this study we used two lines: AB1167, obtained from a healthy donor (herein called ‘healthy’) and AB1071, a line obtained from a DMD patient (herein termed ‘DMD’).

The cell lines were maintained as described in our previous work (Ebrahimi et al., 2021), with the following modifications. For cell expansion prior to tissue seeding, passage 8-10 immortalized myoblasts were plated into 15 cm tissue culture dishes (5×10^5^ cells/dish) within the following growth medium: PromoCell Skeletal Muscle Growth Medium Kit (PromoCell C-23160) supplemented with 15% fetal bovine serum (FBS, Thermo Fisher 12483-020) and 1% Penicillin-Steptomycin (P/S, Gibco 15140-122). Dishes were maintained at 37°C with 5% CO_2_ in a cell culture incubator and half of the culture media was refreshed every other day until reaching 80% confluency, the point at which the cells were used for studies.

### Human iPSC-derived myoblasts and motor neuron production

The iPS11 induced pluripotent stem cell line (Alstem; a kind gift from P. Zandstra) was maintained on Geltrex (Thermo Fisher A1413302; 2%) coated tissue culture treated plates in mTeSR1 medium (STEMCELL Technologies 85850). The cells were split using TrypLE™ Express (Thermo Fisher 12605028) every 3-4 days upon reaching 75-85% confluency.

Myogenic progenitors: iPSC 11 cells were differentiated to the myogenic lineage to produce skeletal muscle myogenic progenitors by following the detailed protocol established by (Xi et al., 2017) with minor modifications to some of the reagents used as detailed here: Geltrex (Thermo Fisher Scientific A1413302; 2%) rather than Matrigel, addition to DNase I (10ug/ml: Millipore Sigma 260913) to collagenase IV and TrypLE during the day 29 dissociation step, and SK Max medium (Wisent Bioproducts, 301-061-CL) as an alternative to SkGM2. Upon establishing a myogenic progenitor line, a portion of passage 0 cells were cryopreserved for future studies. Myogenic progenitors were used for experiments at passages 1 – 3.

Motor neuron clusters: On the first day of differentiation to the post-mitotic motor neuron lineage, the iPSCs were dissociated with TrypLE Express to single cells, 5.8 × 10^5^ cells in N2/B27 media and supplements were seeded into one well of a 6 well AggreWell™400 culture plate (STEMCELL Technologies) which was coated with 5% Pluronic acid F-127 (Sigma-Aldrich P2443) and rinsed with PBS. N2/B27 media consists of DMEM-F12 (Thermo Fisher 11330032) supplemented with N2 (Thermo Fisher 17502048) which is then mixed at a 1:1 ratio with Neurobasal media (Thermo Fisher 21103049) supplemented with B27 supplement (Thermo Fisher 0080085SA). The AggreWell™ plate was then spun at 100 x g for 3 minutes. On the first day of differentiation, the cells in the AggreWell™ plates were cultured in N2/B27 media that was further supplemented with 5 μM Rock-inhibitor (Tocris 1254), 40 μM SB-431542 (Sigma-Aldrich,S4317), 200 nM LDN-193189 (Tocris 6053), 3 μM CHIR99021 (Tocris 4423) and 200 μM ascorbic acid (Sigma-Aldrich, A4403), as described previously (Nijssen et al., 2018) The media was the same on day 2, with the exception that the Rock-inhibitor was excluded. Starting on the third day, the N2/B27 media was instead supplemented with 100 nM retinoic acid (Sigma-Aldrich R2625), 500 nM SAG (Peprotech 9128694) and 200 μM ascorbic acid (Sigma-Aldrich A4403) (Nijssen et al., 2018)and this culture media was used for the remainder of the protocol.

The cell aggregates were maintained in the AggreWell™ plate from day 1 to day 4 with daily media changes as described above. On day 5, the EBs from one AggreWell™ were transferred into two wells of 6-well non-tissue culture treated plate that was pre-treated with 5% Pluronic acid F-127 and the plates were maintained on an orbital shaker at 90rpm until day 10. On day 10, motor neuron clusters were collected for co-culture with the skeletal muscle cells.

### 3D neuromuscular co-cultures

To establish neuromuscular co-cultures, we first fabricated human skeletal muscle macro-tissues using methods similar to our previously published work. Briefly, immortalized human myoblasts or iPSC-derived myogenic progenitors were detached from culture dishes and manually counted. Cells were then spun down and resuspended in an extracellular matrix (ECM)-rich mixture containing 40% DMEM (%v/v, Gibco, 11995073), 40% fibrinogen (10 mg/mL stock concentration in a 0.9% NaCl water solution, Sigma-Aldrich F8630-5G), and 20% Geltrex™ (Thermo Fisher, A1413202) at a density of 8×10^5^ (immortalized) or 1.5×10^6^ cells (iPSC-derived) per 100 μL. Following a 1-hour coating period, pluronic acid (5%, Sigma-Aldrich P2443-250G) was removed from the MyoFMS molds, which were then air dried in a biological safety cabinet for 15 minutes.

Just prior to seeding the MyoFMS molds, thrombin (Sigma-Aldrich. T6884-250UN) was added to the cell-ECM slurry to a achieve a final concentration of 0.2 U thrombin per 1 mg of fibrinogen. After thoroughly triturating the mixture while taking care not to introduce bubbles, 100 μL of the unpolymerized mixture was seeded into each MyoFMS mold and evenly distributed between the micro-post display on either side. After a 10 minute incubation period at 37 °C, the polymerized hydrogel containing human myoblasts was immersed in 2 mL of PromoCell Skeletal Muscle Growth Medium supplemented with 20 % FBS, 1 % antibiotic and 6-aminocaprioc acid at 1.5mg/mL (ACA, Sigma-Aldrich A2504-100G). Polymerized hydrogel containing iPSC-derived myogenic progenitors was immersed in 2 mL of F-10 medium (Wisent, 318-050-CL) with 20% FBS, 1% antibiotic and ACA at 1.5mg/mL. After 2 days (day -2 to 0), growth media was switched to a low-serum differentiating media (DM): DMEM, 2% horse serum (HS, Gibco 16050-122), 1% P/S, 2 mg/mL ACA, and 10 μg/mL human recombinant insulin (Sigma-Aldrich, 91077C).

To produce 3D neuromuscular co-culture tissues, motor neuron clusters were added to muscle tissues on day 0 of differentiation. More specifically, iPSC-derived motor neuron clusters were detached from their culture substrate and spun down. Afterwards, the fibrin/Geltrex master mix described above was prepared and then, using a p200 micropipette, 5-6 motor neuron clusters (∼300 - 400 microns in diameter) were added for every 25 μL of ECM master mix. The day 0 muscle macro-tissues were removed from the incubator, placed within a biological safety cabinet, and the media was completely aspirated. Thrombin (0.2 U/mg) was incorporated into the slurry and 25 μL (including 5-6 neuron clusters) was then added dropwise to each side of a MyoFMS macro-tissue. After a 15-20 minute gelation period at 37°C, neuromuscular co-cultured were gently covered with 2 mL of DM containing brain-derived neurotrophic factor (BDNF, 10 ng/mL, PeproTech 450-02-10μg) and glial cell-derived neurotrophic factor (GDNF, 10 ng/mL, PeproTech 450-10-10μg). Half media changes were conducted every other day, which included addition of the neurotrophic factors at a two-fold concentration.

### Electron microscopy sample preparation and imaging

Tissue embedding and processing was previously described in Nguyen et al., 2021. In brief, tissues were fixed in 2.5 % glutaraldehyde and 2 % paraformaldehyde in 0.1 M sodium cacodylate buffer for 2 hours at room temperature. Sample embedding and sectioning was completed at the Nanoscale Biomedical Imaging Facility at The Hospital for Sick Children, Toronto Canada. Muscle tissue samples were washed in 0.1 M sodium cacodylate buffer with 0.2 M sucrose and fixed for 1.5 hours on ice in 1 % osmium tetroxide and washed in cacodylate buffer. An alcohol series of dehydration occurred where muscle samples were washed in 50%, 70%, and 90% ethanol for 20 minutes each and then washed three times in 100 % ethanol.

Ethanol was displaced by 100% propylene oxide twice for 40 minutes. Muscle samples were then placed in 50:50 propylene oxide/Quetol-Spurr resin for 2 hours, then infiltrated in 100 % resin overnight. Muscle samples were then embedded in resin mould and baked overnight at 65 ºC. Muscle samples were then sectioned longitudinally, collected on mesh grids, and stained with lead citrate and uranyl acetate. Images were processed and collected on a FEI Technai 20 electron microscope.

## Results

Here we present our results obtained from three different muscle types, grown in the presence or absence of co-cultured iPSC-derived MN clusters. Macrotissues were grown from i) immortalized healthy muscle cells, ii) immortalized DMD muscle cells, and iii) iPSC muscle cultures. Our analysis confirms our previous observations that tissues derived from DMD donors are disorganized compared to tissues grown from healthy donors. We further add that the presence of co-cultured MNs substantially improves myotube ultrastructural organization of the contractile apparatus. By providing a qualitative assessment of muscle structural features of the sarcomere Z-discs, M-lines, I and A bands, and H zones, we report a positive effect of motor neuron presence on the sarcomere organization of 3D cultured muscle. Previously, we described a method to generate human neuromuscular co-cultures in a custom device consisting of a seeding channel with Velcro™ attachments at each distal end (Bakooshli et al., 2019). Unfortunately, we found that about half of all neuromuscular co-cultures detached from the Velcro™ hooks within a week, rendering them unusable for analysis. Anecdotally, innervated muscle tissues produced very strong spontaneous contractions, and we hypothesized that the original culture platform was unable to support the contractions. Therefore, we reimagined the design, and herein report a new co-culture device, which we refer to as MyoFMS (**myo**tube **f**unction in **m**ulticellular **s**caffolds; Figure 1), which eliminated the loss of neuromuscular co-cultures observed in the prior device.

**Figure 1.**
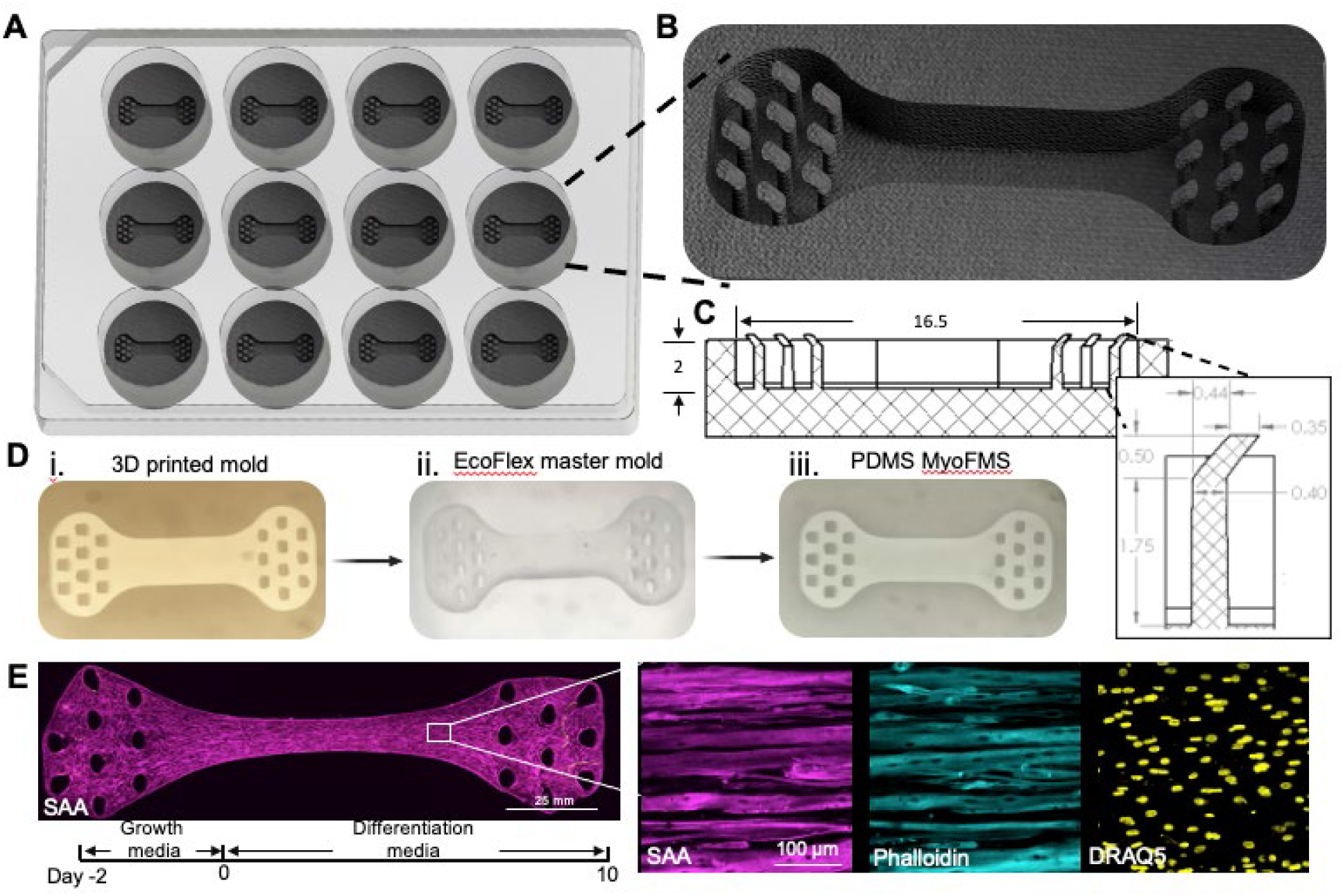
MyoFMS device fabrication and skeletal muscle tissue production. (**A**) 3D model of a 12 well foot-print MyoFMS culture plate together with a (**B**) 3D model zoom-in and a (**C**) CAD drawing of a single well to visualize multi-post arrangement and tissue seeding bed dimensions and orientation. (**D**) Representative images of each step in the fabrication process used to generate the final PDMS-based MyoFMS culture device. (**E**) Timeline showing the steps employed to generate skeletal muscle tissues in MyoFMS. (**F, left**) Representative, 4x stitched confocal image of iPSC-derived skeletal muscle tissue at day 10 of differentiation. Magenta, sarcomeric α-actinin (SAA). Scale bar, 25 mm. (**F, right**) 40x confocal split-channel images of a representative region of the (**F, left**) engineered muscle tissue. Magenta, SAA; Cyan, phalloidin; yellow, Draq5. Scale bar, 100 μm.

### Tissues from Immortalized Healthy Cells

Muscle fibers of healthy tissues in both the presence (*n*=2) or absence (*n*=3) of iPSC-derived MNs exhibit myofilament sarcomere organization typical for human muscle. Alternating A-I bands are observed in both the muscle alone and nerve-muscle co-cultures (Figure 2). There is some variation in the degree of sarcomere banding and organization between single muscle fibers within the same tissue. In contrast, tissues co-cultured with iPSC-derived MNs uniformly possess a high degree of distinct myofibrils and filament banding. Additionally, sarcomere organization improved in all muscle fibers, resulting in clearer A-I banding, and well-defined sarcomeres compared to muscle-alone cultures.

**Figure 2.**
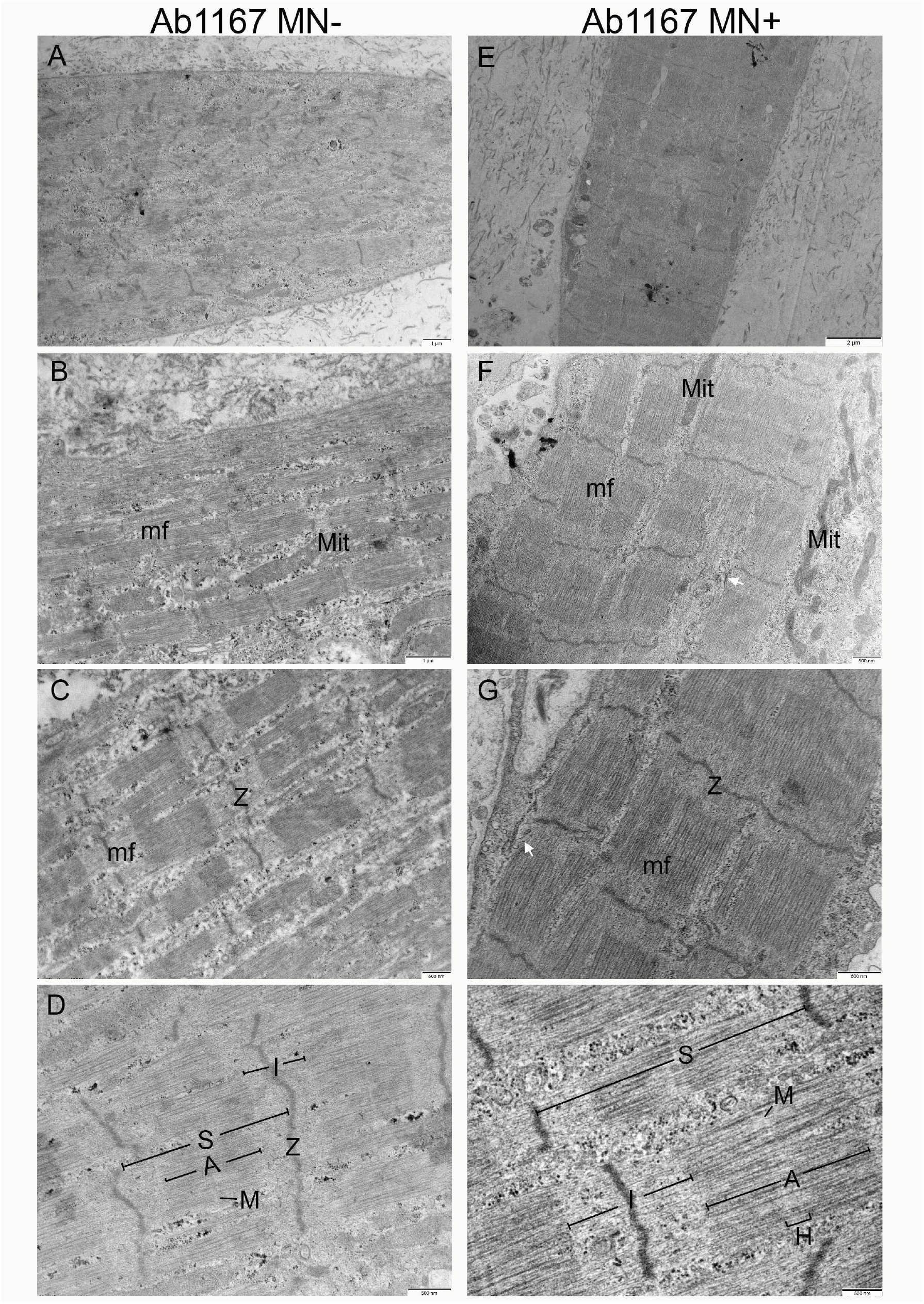
Representative comparison electron micrographs of 3D bioengineered macrotissues +/-iPSC-derived motor neurons at Day 10. **(A-D)** Healthy AB1167 muscle-alone (*n*=3), **(E-H)** neuro-muscular AB1167 co-cultured with iPSC-derived motor neurons (*n*=2). **(A and E)** Single muscle fiber showing multiple myofibrils (mf). Ultrastructural features in both conditions include mitochondria (Mit), sarcomeres (S), A-bands (A), I-bands (I), H-zones (H), M-lines (M), and Z-discs (Z). (A-D) macrotissues show distinct myofibrils with clear myofilament organization. **(E-H)** neuro-muscular macrotissues show slight improvement in sarcomere organization with clearer M-lines, and strong crosslinking compared to muscle-alone tissues. Sarcoplasmic reticulum (SR) becomes more prevalent in neuro-muscular macrotissues (white arrow).

**Figure 3.**
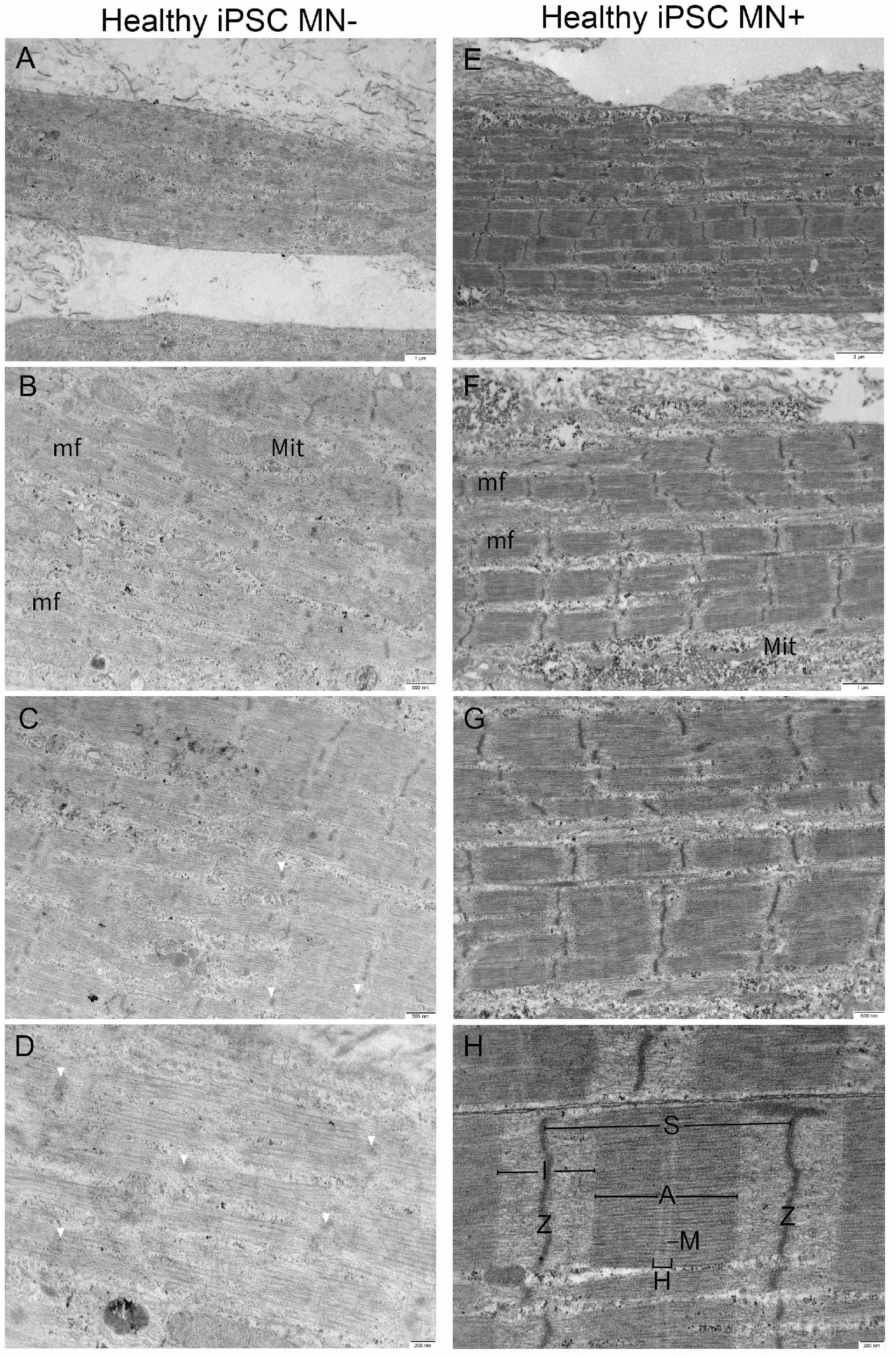
Representative comparison electron micrographs of 3D bioengineered macrotissues +/-iPSC-derived motor neurons at Day 10. **(A-D)** Healthy iPSC-derived muscle-alone (n=1), **(E-H)** neuro-muscular iPSC derived muscle co-cultured with iPSC-derived motor neurons (n=1). **(A and E)** Single muscle fiber showing multiple myofibrils (mf). **(E-H)** Mitochondria (Mit) is seen in both conditions, whereas faint short Z-disc (Z), less distinct sarcomere (S), and structural features A-bands (A), I bands (I), H-zones (H), and M-lines (M) is seen in muscle-alone tissues **(A-D)** compared to neuro-muscular tissues **(E-H)**. In the presence of motor neurons myofilament organization improves, illustrated with clear Z-disc, A-I banding, strong crosslinking with well-defined M-line and H-zones.

We also observe presumptive sarcoplasmic reticulum (SR) and T-tubules (TT) profiles in both muscle-alone and co-cultures, but these become increasingly prevalent and more defined in MN co-cultures (Figure 5 (D and E)). SR-TT triad structures were observed in myofibrils in MN co-cultures (Fig. 5 (A and B)), whereas they were not seen in muscle-alone cultures. While the boundaries of these triads may not have the same clarity as seen in electron micrographs of biopsied human muscle, their close proximity to each other paired with dense membranes suggests a triad.

We measured the sarcomere length from a subset of these tissues and found sarcomeres were 2.05±0.02 μm in the healthy muscle alone and 1.87±0.03 μm in healthy muscle plus MNs. A nested T-test of the data obtained from 3 independent tissues of each type showed the difference between the means was not significant.

### Tissues from Immortalized DMD Cells

The majority of DMD muscle fibers, in the absence of MNs, (*n*=3) were disorganized and only 10% of myotubes had some degree of skeletal muscle banding. Muscle-alone fibers consisted of a dispersed network of myofilaments in non-uniform directions across the entire fiber, making individual myofibrils indistinguishable (Fig. 4 (A-D)). While some myofilaments were grouped together forming dense sarcomere silhouettes, they lacked the distinct features usually present in a complete sarcomere structure as shown in Fig. 4 (B and C). Additionally, Z-discs appeared not fully formed but rather as electron-dense fragments that could be seen throughout these fibers (Figure 4D).

**Figure 4.**
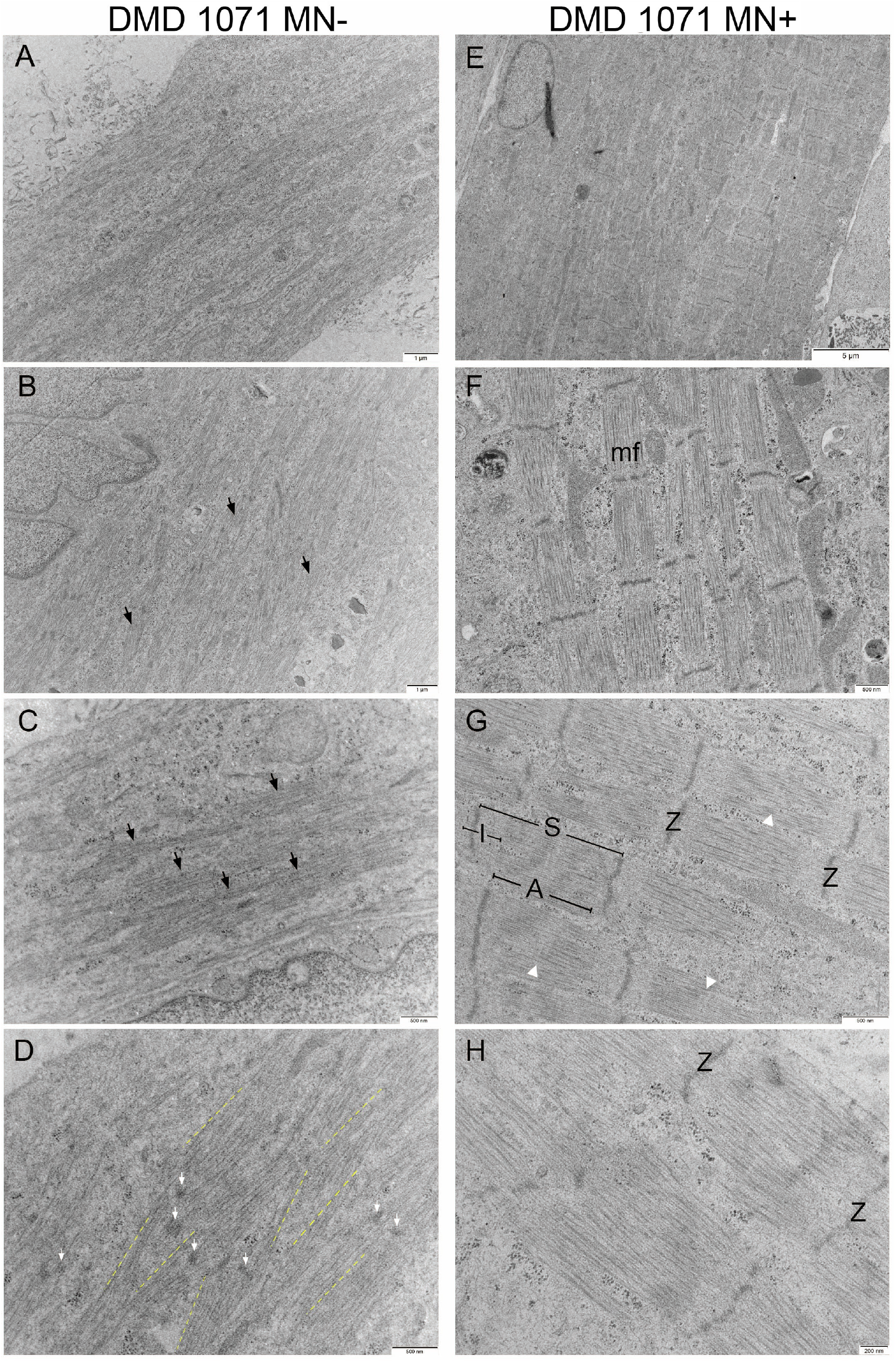
Representative comparison electron micrographs of 3D bioengineered DMD-1071 macrotissues +/-iPSC-derived motor neurons at Day 10. **(A-D)** DMD 1071 muscle-alone (*n*=3), **(E-H)** DMD 1071 co-cultured with iPSC-derived motor neurons (*n*=4). **(A and E)** Single muscle fiber. **(A-E)** DMD 1071 muscle-alone electron micrographs showing the presence of disorganized myofilaments (black arrows). (**D**) Myofilaments run in non-uniform, unparalleled directions (yellow dashed lines). Faint and short z-bodies are seen throughout tissues with ill-defined sarcomere boundaries (white arrows). **(E-H)** DMD 1071 neuro-muscular macrotissues show a significant improvement in sarcomere organization showing the presence of sarcomeres (S), A-bands (A), I bands (I), and Z-discs (Z), and some faint M-lines (white arrowhead), when compared to muscle-alone.

In contrast to DMD muscle-alone cultures, DMD muscles in the presence of iPSC-derived MNs (*n*=4) appeared to show an improvement in organization. DMD muscle fibers with MN co-cultures consistently possess distinct myofibrils with clear sarcomeres and Z-discs.

Although thick and thin filaments making up the A- and I-band are not as prominent and dense compared to healthy fibres, the M-lines and H-zones can still be identified, albeit faintly (Fig. 4G). Similar to healthy tissues, presumptive SR-TT profiles were rarely observed in the muscle-alone fibers, however become more well-defined and prevalent across the muscle fibers and in between myofibrils of DMD co-cultures (Fig. 5F). No muscle triads were observed in DMD macrotissues. Though sarcomere organization significantly improves in the co-cultures with identifiable myofibrils and sarcomeres, when compared to healthy tissues, the high incidence of ill-defined M-lines and H-zones and less dense A and I bands indicated that the DMD co-cultures were less well developed (Figure 4 (G and H); Table 1).

**Table 1.** Grading the characteristic presence of contractile skeletal muscle features found in human 3D bioengineered macrotissues +/-iPSC-derived motor neurons.

|  | Healthy |  |  |  | DMD |  |
| --- | --- | --- | --- | --- | --- | --- |
|  | AB1167<br>MN-<br>(n=3) | AB1167<br>MN+<br>(n=2) | iPSC<br>MN-<br>(n=1) | iPSC<br>MN+<br>(n=1) | DMD 1071<br>MN-<br>(n=3) | DMD 1071<br>MN+<br>(n=4) |
| *Myofibrils* | ••• | •••• | •• | •••• | • | •••• |
| *Sarcomere* | ••• | •••• | •• | •••• | • | ••• |
| *Z-disks* | ••• | •••• | •• | •••• | • | ••• |
| *A-band* | ••• | •••• | •• | •••• | • | ••• |
| *I-band* | ••• | •••• | • | •••• | • | ••• |
| *M-line* | ••• | ••• | • | ••• | • | •• |
| *H-zone* | ••• | ••• | •• | ••• | •• | ••• |

\*n = number of macrotissues examined per myogenic line
\*\* 3D bioengineered macrotissues are compared to a perfect score of 5 dots, representative of a human muscle biopsy.

**Figure 5.**
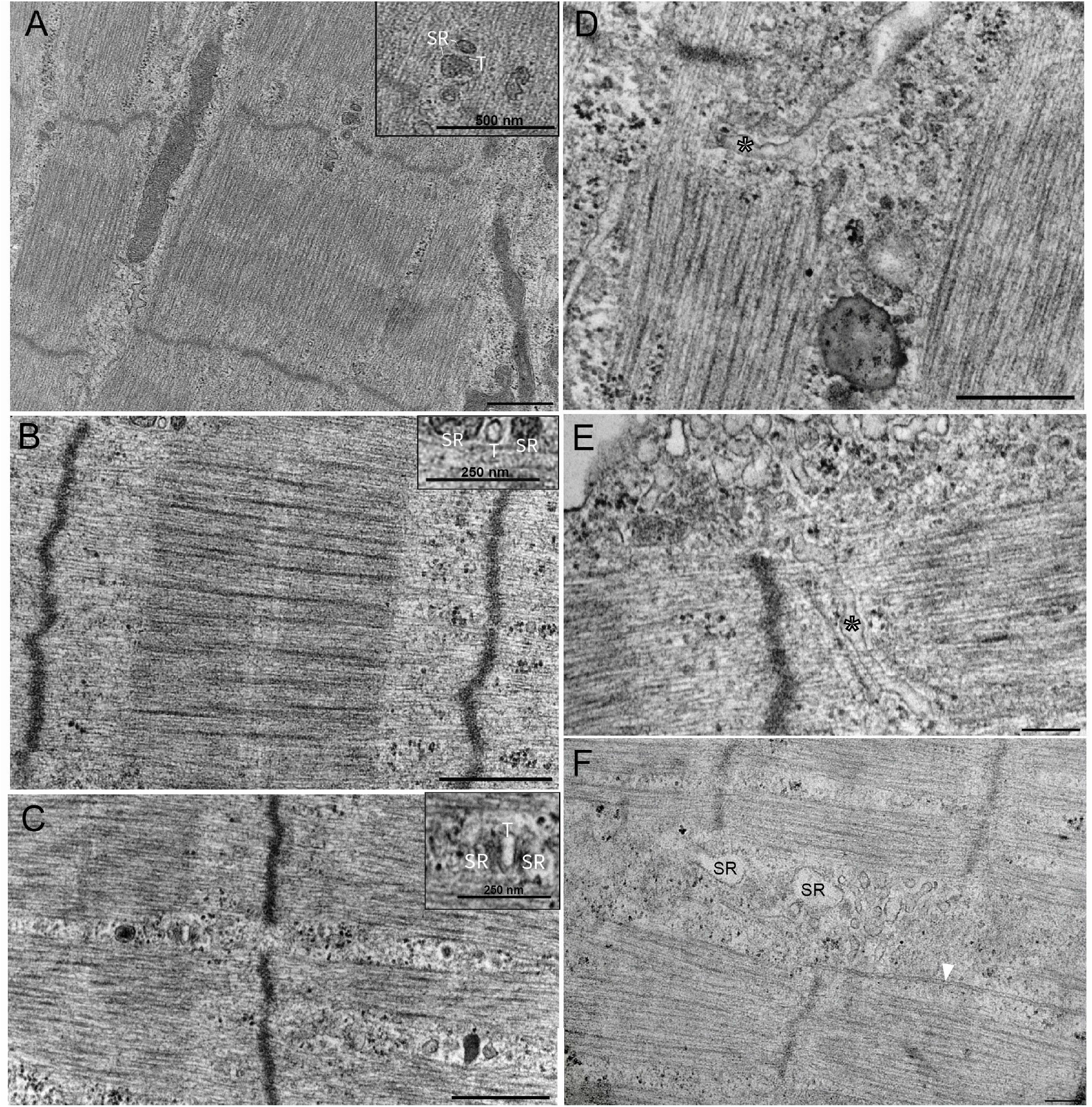
Ultrastructural features of 3D bioengineered neuromuscular macrotissues at Day 10. **(A and B)** Healthy Ab1167 co-culture illustrating presumptive skeletal muscle triad with sarcoplasmic reticulum (SR) and t-tubule (T) between myofibril. Scale 500 nm. **(C)** Healthy iPSC co-culture illustrating presumptive skeletal muscle triad with sarcoplasmic reticulum (SR) and t-tubule (T) between A-I band. Scale 500 nm. **(D and E)** Healthy Ab1167 co-culture illustrating t-tubule invagination through myofibril (asterisk). Scale 500 nm. **(F)** DMD 1071 co-culture showing sarcoplasmic reticulum (SR) and microtubule (white arrowhead) between myofibrils. Scale 200 nm.

### Muscle Tissues derived from iPSCs

We show for the first-time ultrastructure of 3D cultured muscle tissues derived from iPSCs (*n*=1), as well as robust and functional iPSC neuro-muscular co-cultures (n=1). Using a healthy PSC line (ALTSTEM iPSC11; male) (Xi et al. 2017; Hicks et al. 2018), we identified an optimum cell seeding density to generate iPSC macrotissues populated by multinucleated myotubes, along with incorporating motor neurons derived from iPSCs to create our co-culture system. Consistent with our immortalized healthy and DMD macrotissues, myotubes from healthy iPSC co-cultures had improved sarcomere organization compared to muscle alone cultures (Fig.3).

The myotubes in iPSC muscles-alone showed some variability in structure. Most of the muscle fibers possessed recognizable sarcomere elements, some sarcomeres varied in shape and size, as illustrated in Fig.3. A small number of fibres in the sample were disorganized, with indistinguishable or unorganized myofilaments with ill-defined structural features such as M-line and H-zones.

The iPSC derived muscle co-cultured with iPSC derived motor neurons showed dramatically improved myofibril and sarcomere organization, resulting in clearly defined A-I banding, M-lines, H-zones, and Z-discs (Table 1). SR-TT profiles were apparent in both the muscle-alone and MN co-culture conditions. Whereas the SR-TTs are randomly dispersed across the muscle fiber in muscle-alone, they become more localized to the gaps between myofibrils in the MN co-cultured tissues. Furthermore, there appears to be a higher amount of the SR-TT structures in the iPSC muscle-alone and muscle co-cultures, in comparison to the tissues derived from the immortalized cells lines. SR-TT triad structures can be observed in myofibrils in healthy co-cultures (Fig. 5C).

## Discussion

The central advance of this report is that in three different muscle types grown in 3D culture, healthy and DMD immortalized cell lines and muscle derived from iPSCs, motor neuron co-culture improves the organizational appearance of the sarcomeres. In the presence of MNs, myofibrils appear to have better thick and thin filament alignment, more prominent A and I bands, continuous and aligned Z-discs, and more obvious presence of SR and TT profiles that come together as triads.

The findings herein are noteworthy for several reasons. This new data confirms our previously published work (Ebrahimi et al., 2021; Nguyen et al., 2021), that muscle-alone cultures of DMD derived tissues are disorganized compared to those derived from healthy donors. The results in this paper, together with those in Nguyen et al. 2021 and Ebrahimi et al 2021 now give us three independent examinations of muscle-alone cultures representing a total of 8 independently cultured DMD tissues examined across all studies and the data uniformly show disorganization in DMD tissue. This observation was fundamentally enabled by our culturing platform and the ability to separate muscle from nerve, which is not possible for tissue obtained via biopsy.

The finding that MNs exert a positive influence on both DMD and healthy tissues suggests several lines of investigations which may lead to new therapeutic strategies that harness this natural nerve-muscle support. For example, at present we do not know whether synaptic contacts are necessary for this organizational improvement or whether the MN clusters are secreting trophic substances which aid the muscle. Our prior studies (Bakooshli et al. 2019) have already confirmed functional synaptic contacts with our culture methods and though we have ongoing studies to describe synaptic ultrastructure, classical cell-separation or conditioned-media assays may be helpful to delineate these two possibilities and suggest further investigations.

While the neurotrophic relations (Guth, 1968; Gutmann, 1976) between nerve and muscle have been well established, clues regarding the trophic support of muscle by neurons may be gained from studies such as (Saini et al., 2021) who identified several trophic factors that appeared to be upregulated in muscle-nerve co-cultures. It remains to be determined whether trophic factors released from MNs directly support muscle organization or alternatively, the presence of MNs and/or synaptic connections induces muscle to produce and secrete trophic factors that act in a paracrine manner. In either case, once identified, such factors may prove useful to ensuring better ultrastructural organization of DMD muscle.

The ultrastructure of dystrophic muscle has long been the subject of electron microscopic studies and indeed EM has been a key technology to advance important ideas about DMD pathology, such as the membrane leakage hypothesis (Mokri & Engel, 1975). Several early examinations of ultrastructure suggested that there were no observable differences in the myofibrils between healthy and dystrophic muscle (Shafiq et al., 1969; Wechsler, 1966) but the authors concede the limited nature of these conclusions since at the time, very little was known about the cause or progression of disease. At least one early study did identify myofibril damage in preclinical cases of DMD (Pearce, 1966).

It is generally acknowledged that 2D muscle cultures do not replicate the complex sarcomeric architecture of *in vivo* muscle (Dessauge et al., 2021). While cultures of C_2_C_12_ mouse muscle cells and more recently human muscle cells have been studied and show muscle specific protein expression (Burattini et al., 2004; Owens et al., 2013) they do not develop the typical banding patterns of normal muscles observed at the level of light or electron microscopy. Better ultrastructure is seen when C_2_C_12_ cells grown in collagen-based 3D matrix (Rhim et al., 2007), in which cells did show striations at both the light and EM level, in contrast to the 2D cultures.

Interestingly, study of muscles cells grown from iPSCs in 2D culture (Skoglunda et al., 2014) also indicated better sarcomere organization than other reports of 2D culture. Together, with our report of iPSCs in 3D culture, and the particularly robust organization of sarcomeres in the presence of MNs, we suggest that tissues grown from iPSCs have the potential for the best developmental profile of muscle cells in culture so far reported. Whereas the iPSCs in 2D culture developed good sarcomere organization in about 20 days (Skoglunda et al., 2014), the results shown here were obtained in 10 days, further indicating the advantages of the 3D culture platform.

In sum, our study provides compelling evidence that motor nerves contribute to the process of sarcomere development, and suggest that in dystrophin-deficient muscle cells, may serve to overcome deficits in the earlier steps of myofibrillar growth. The modular and tractable nature of the approach described herein offers an ideal setting with which to explore the cellular and molecular mechanisms driving these effects, which in turn offers the possibility of new therapeutic interventions.

## Acknowledgements

The study was supported by the Canadian Institutes of Health Research Project Grant #PJT-168932 (to B.A.S. and P.M.G.), Ontario Institute for Regenerative Medicine Grant 2018-0510 (to P.M.G.), Canada First Research Excellence Fund Grant MbDC2-2019-02 “Medicine by Design” (to P.M.G.), and Canada Research Chair in Endogenous Repair Award 950–231201 (to P.M.G.). We thank Ali Darbandi at the Nanoscale Biomedical Imaging Facility at The Hospital for Sick Children, Toronto for assistance with transmission electron microscopy.

